# Fighting Through the Heat: How Sexual Selection Influences Demography Under Recurrent Heatwaves

**DOI:** 10.1101/2025.03.31.646346

**Authors:** Neelam Porwal, Jonathan M. Parrett, Agnieszka Szubert-Kruszyńska, Neha Pandey, Robert J. Knell, Tom C Cameron, Jacek Radwan

## Abstract

Sexual selection is a potent evolutionary force with complex effects. Strong sexual selection can enhance adaptation and reduce mutational load, while simultaneously reducing survival, or causing sexual conflict that reduces fitness for one or both sexes. Many populations today face not only gradual environmental changes but also extreme, short-term stress events like droughts or heatwaves. The combined effects of sexual and environmental selection on population demography during and after such events remain poorly understood, even though such combined effects could be crucial for the persistence of small, endangered populations under climate change. In this study, we investigated how sexual selection affects survival during environmental stress by manipulating the expression of an aggressive fighter morph in small populations of the male-dimorphic soil mite *Sancassania berlesei,* exposing some of these populations to recurrent periods of extreme heat and monitoring survival over eight generations. We found that heat exposure reduced survival, more severely in females than in males, and survival was lower in populations with higher fighter prevalence, but there was no interaction between temperature and fighter morph prevalence. Furthermore, survival declined across generations, and the decline was steeper in populations with lower prevalence of fighters, leading to the loss of their initial survival advantage by the last generation. T265625hree populations exposed to heat became extinct during the experiment, all from the reduced fighter expression treatment. Our findings imply that despite its cost to individual survival, sexual selection does not modulate population sensitivity to heatwaves over several generations. Furthermore, we demonstrate that these mortality costs of sexual selection are gradually compensated over successive generations, which could be a result of a more effective purging of inbreeding depression. Thus, while the additive effect of sexual selection and heatwaves on survival may increase demographic risks for bottlenecked populations in the short term, sexual selection may increase resilience of populations to prolonged bottlenecks.

## Introduction

Global environmental change, including rising temperatures, present substantial risks to biodiversity (Habibullah et al., 2022; Ripple et al., 2017), with notable declines forecast for species lacking adaptive capacity or dispersal abilities even with a modest increase of 1–2 °C in temperature (Nunez et al., 2019). One of the most important predicted consequences of climate change is increased variability of climate, and a crucial aspect of this is likely to be an increase in the frequency and the amplitude of heatwaves. Heatwaves, are known to have stronger impacts on life than directional changes (Sheldon & Dillon, 2016) by directly influencing individual survival, physiological processes, and reproductive efficacy (Boni, 2019) with these effects being more pronounced in ectothermic species (Kingsolver et al., 2013; Ma et al., 2021; Neven, 2000; Walsh et al., 2019). Under such conditions populations may be selected to evolve towards enhanced thermal tolerance, with traits that may enhance survival in high-temperature environments being favoured, potentially allowing “evolutionary rescue” to occur (Bell, 2017). However, adaptation may be severely limited when populations decline to the size limiting the amount of genetic variance available for selection (Frankham, 2010). Furthermore, small population sizes result in significant genetic drift and inbreeding depression (Spielman et al., 2004). This when combined with environmental stressors, can create a harmful cycle referred to as the “vortex of extinction” that can elevate the risk of extinction (Godwin et al., 2020; Ivimey-Cook et al., 2021; Reed et al., 2012; Soulé et al., 1986).

Understanding how populations adapt before environmental change drives them to extinction is a key research focus (Hoffmann & Flatt, 2022; McCulloch & Waters, 2023; Nunney, 2016). One mechanism that may influence adaptability is sexual selection with multiple processes identified that drive the co-evolution of thermal ecology and sexually selected traits. These processes illuminate the role of sexual selection in facilitating thermal adaptation. Anthropogenic warming affects sexual selection, which can either promote adaptation by eliminating harmful alleles or hinder it through inbreeding and the evolution of costly traits (Lieth et al., 2022). As a result, temperature stress can alter sexual selection dynamics, potentially impacting population persistence (García-Roa et al., 2020).

Sexual selection is a process arising from competition for mates and their gametes that favours traits which bestow competitive advantages in the context of mating, such as intricate displays serving to attract mates, armaments and behaviours useful in direct competition for them or seminal fluids aiding competition for gametes (Andersson, 1994; Darwin, 1871). Additionally, traits used in reproductive competition are costly and therefore only the males in good condition, reflecting their adequate adaptation and/or low mutation load should achieve higher reproductive success (Zahavi, 1979; Rowe and Houle, 1996). This could align sexual selection with natural selection, resulting in positive effects of the sexual selection on population viability (Candolin & Heuschele, 2008; Lorch et al., 2003).

Sexually selected traits may, however, decrease survival of their bearers (Ditchkoff et al., 2001; Endler, 1980), or cause sexual conflict via their negative side-effects on female fitness, directly, e.g. when coercive copulations inflict harm on females (Parker, 1979; Rowe et al., 1994) or indirectly, via negative pleiotropic effect on female fitness (Harano et al., 2010; Rice & Chippindale, 2001), potentially leading to net negative effects on population viability (Flintham et al., 2023). Strong skew in reproductive success resulting from sexual competition or skew in sex ratio due to male harm on females may further amplify these negative consequences by reducing effective population size (Grayson et al., 2014), increasing drift and inbreeding at the cost of adaptability. All these possibilities imply that sexual selection may be of key significance for populations endangered with extinction, particularly under environmental challenge (Kokko & Brooks, 2003). The costs of exaggerated sexually selected traits and associated sexual conflicts (García-Roa et al., 2019; Parsons, 1995; Riechert, 1988), combined with the negative consequences for survival of environmentally challenged populations discussed above possibly are further augmented by demographic stochasticity (Martínez-Ruiz & Knell, 2017). Sexual selection can influence both the immediate effects of environmental stress on individual fertility and survival (Baur et al., 2024; García-Roa et al., 2020; Moiron et al., 2022), and long-term adaptation (Gómez-Llano et al., 2024; Iglesias-Carrasco et al., 2024; Parrett & Knell, 2018; Plesnar-Bielak et al., 2012).

Studies on the effects of sexual selection on population viability so far have reported variable outcomes. Positive effects of sexual selection are well supported in experimental evolution studies (reviewed in Cally et al., 2019), with both purging of mutational load in stable environments (e.g. (Lumley et al., 2015; Parrett et al., 2022) and enhanced adaptation to novel environments (e.g. Plesnar-Bielak et al., 2012) being demonstrated. However, other sets of laboratory studies have reported significant negative effects of sexual selection via sexual conflict (Berger et al., 2016; Holland & Rice, 1999). Additionally, comparative studies have linked elaboration of male sexually selected traits to increased risk of extinction (Bro-Jørgensen, 2014; Martins et al., 2018; but see Morrow & Fricke, 2004; Morrow & Pitcher, 2003), although other studies reported that the reverse relationship in populations subjected to anthropogenic disturbances (Moore et al., 2024; Parrett et al., 2019), suggesting interactive effects of environmental factors.

Experimental evolution studies allowing manipulation of both sexual selection and environmental stress are well suited to examine such interactive effects. Most studies to date have manipulated sexual selection by either manipulating its intensity by altering adult sex ratios or manipulating mating systems either allowing polyandry or enforcing monogamy. Such manipulation either during (Plesnar-Bielak et al., 2012) or prior to (Berger & Liljestrand-Rönn, 2024; Godwin et al., 2020; Iglesias-Carrasco et al., 2024) environmental challenge generally showed positive effects of sexual selection, particularly in environments limiting the possibility of female harassment by males (Berger & Liljestrand-Rönn, 2024; Yun et al., 2018). However, to what extent these results can shed light on the conflicting results discussed above is unclear. Most of these studies focused on step or gradual environmental change, whereas extreme weather anomalies, such as heat waves, may dominate environmental effects on demographics of natural populations, as discussed above.

Furthermore, these studies mostly focus on consequences of manipulation of ecological processes (e.g. via sex ratio manipulation), which by themselves can alter the demographic parameters we would be measuring on a population scale. Indeed, one study which did manipulate sexual selection by using evolutionary lines artificially selected for or against the presence of armoured aggressive males found contrasting results. In *Rhizoglyphus robini* exhibiting alternative reproductive phenotypes, lines nearly fixed for costly weapons were more likely to go extinct under gradual temperature increase (Łukasiewicz et al., 2023).

We conducted a multi-generational experiment to examine how male morph ratios influence population demography under heatwaves using another acarid mite *Sancassania berlesei.* This species exhibits similar male dimorphism to *R. robini*, but with morph heritability being low and mostly determined by population density during nymphal development (Radwan, 1995). As the cue to the density is provided by pheromones (Radwan et al., 2002), we can manipulate the expression of fighter morphs in populations independently of genetic background, allowing us to investigate the effect of manipulation of proportions of sexually selected males on population demographics in the context of thermal stress. We contrasted populations with a higher proportion of fighter males (pheromone control) against those with a lower proportion of fighter males (pheromone treatment). Within each pheromone regime, we either maintained populations under a stable temperature environment or subjected them to successive periods of extreme heat over eight generations to simulate the thermal stressors many species may encounter due to climate change. Importantly, this kind of an intermittent stress may have only passing effects on individual condition, and thus not suppress expression of costly sexually selected traits. This is a relevant point because by reducing associated costs, such suppression by chronic stress (e.g. associated with permanent environmental change) was hypothesised to alleviate negative effects of sexual selection on population viability (Kokko & Brooks, 2003). To ensure expression of fighter morph is not suppressed by stress in our study, we applied it at the adult stage when the morphs are fixed.

We predicted that high prevalence of fighter males will have a negative effect on the survival of males due to fights, but possibly also of females which are also sometimes killed by fighter males (Łukasik, 2010). We further predicted that heatwaves (defined as 45 hrs of extreme rise in temperatures in our experiment) would increase mortality and may have disproportionately more detrimental effects on fighters because their expression of costly weapons could make them less tolerant to additional costs imposed by thermal stress.

## Methods

### Housing and Maintenance of Mite Populations

For this study, we used soil mites (*Sancassania berlesei*), originally collected from poultry litter from a farm in Dluga Goslina, Wielkopolska region of Poland, in 2022. These populations have since been maintained under controlled conditions, with overlapping generations and large population sizes (>1000 adults). Populations were housed in two custom-designed bottle-top enclosures consisting of the bottleneck and cap with a 5mm hole tightly sealed with a cotton plug, and a cemented base, to balance ventilation and containment. Maintenance conditions included high humidity levels (over 90%) and a constant temperature of 23°C in dark incubators. The mites were fed dry yeast twice weekly and were transferred to new bottle tops and mixed between two bottle-tops once monthly.

### Experimental populations

To obtain generation-synchronised experimental populations, adult mites from the stock populations were isolated into a new box for a 2-day egg-laying period. After the eggs hatched (4 days after adults were removed), 50 larvae were transferred to each of 42 glass vials (2.5 cm x 5 cm). The bottom of each vial was covered with a 2.5 cm layer of plaster of paris to create a water-absorbent substrate. The vial lids had a 10 mm hole covered with fine mesh to facilitate air exchange while preventing mite escape. To minimize disturbance, excess food was provided but only during vial transfers. Populations were assigned to one of two treatments: pheromone treatment was used as described below to decrease the proportion of fighters in a population compared to pheromone control populations. Both these treatments were split into two temperature treatments (heatwave vs. stable) in a full 2×2 factorial design consisting of 15 pheromone treatment heatwave, 15 pheromone control heatwave ones, 6 pheromone treatment stable and 6 pheromone control stable populations.

We created more heat-exposure compared to control populations because we expected heat exposure to lead to population extinction, reducing the number of replicates.

### Pheromone Treatment

Populations from both pheromone regimes, during their development from larvae to adults (pheromonal sensitive window of morph determination), were maintained in the pheromone treatment setup. This setup consisted of two transparent plastic boxes; one placed atop the other, separated by a lid. The lid of the bottom box and the base of the top box had square cutouts covered with mesh to allow pheromone diffusion between the boxes. Bottom boxes contained two high-density mite populations housed in plastic containers. While these populations were not counted, we estimate each contained hundreds of thousands of mites. Each container had a 1 cm cement base and was filled with a ∼2 cm layer comprising a 1:1 ratio of food to mites. The containers were surrounded by dishwashing liquid to prevent mites from contaminating experimental populations. The pheromone treatment populations were placed in the top box, which was covered with a lid featuring holes covered with porous tape to permit restricted air exchange (Fig.1). Concurrently, both pheromone controls were maintained in an identical setup without the high-density mite populations to ensure consistent environmental conditions. Logistically it was only possible to set up one container each for pheromone control and pheromone treatment and then transferred to different temperature treatments respectively, but the containers hosting both treatments were identical except for the presence of dense colonies in pheromone treatment, and given otherwise highly standardised conditions we do not expect any uncontrolled difference between containers to affect our results.

**Fig. 1.**
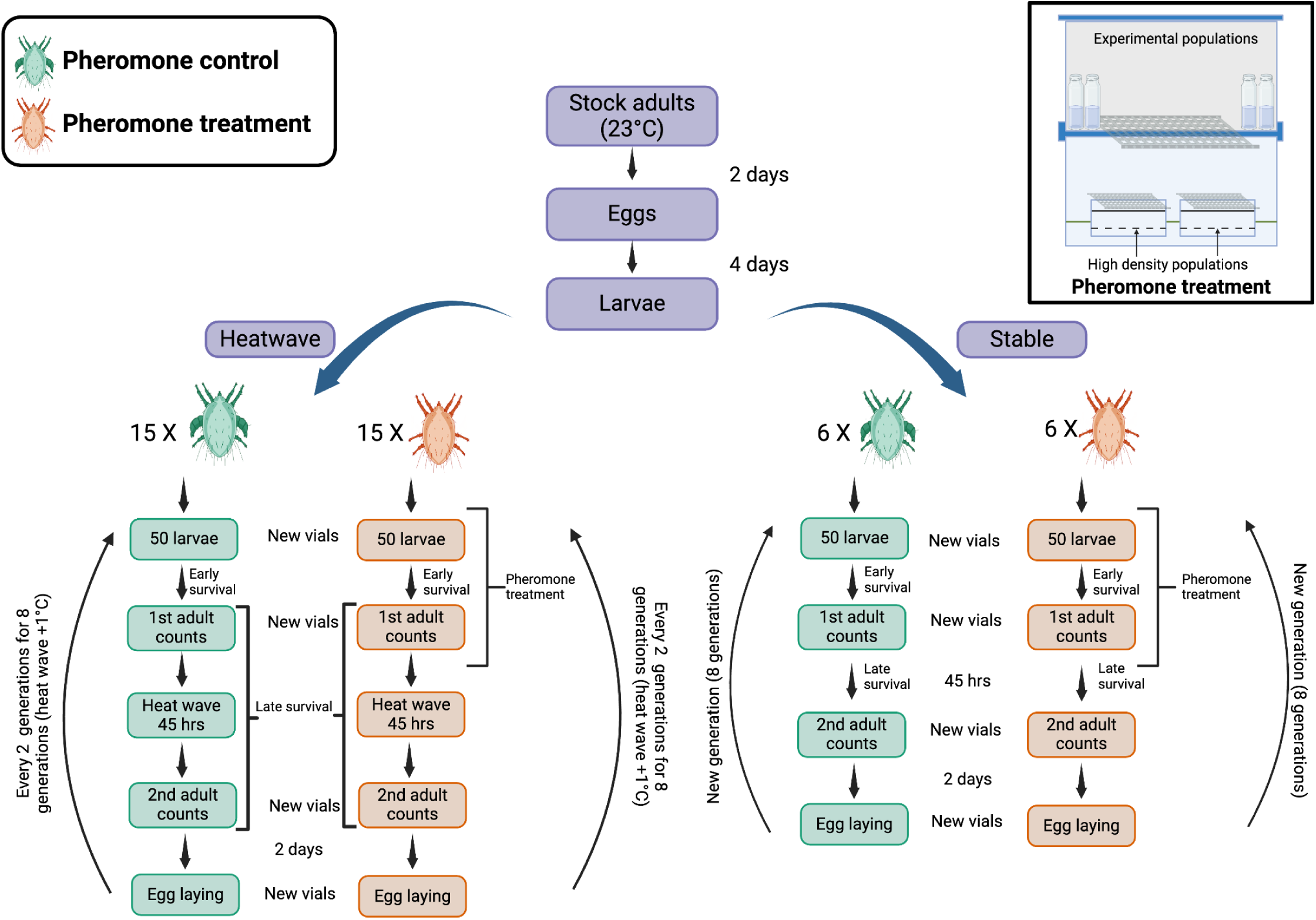
Schematic of the experimental workflow, illustrating the pheromone control (green) and pheromone treatment (orange) regimes. Stock adults (23°C) produced eggs that developed into larvae over four days before being assigned to either heatwave or stable environments. Pheromone treatment populations were exposed to high-density pheromone cues from the larval stage until the first adult counts in the setup on the top right. Key experimental steps, including survival assessments, developmental stages, and reproductive phases, are outlined for both conditions. Created in https://BioRender.com

### Experimental Workflow

Larvae from the F_0_ generation were transferred into 42 vials and randomly assigned to one of two treatment groups: pheromone exposure (treatment) or no pheromone exposure (control). Vials were maintained at 23°C, and treatments were applied for a period of 6 to 8 days. First adult counts were conducted 6 days post-transfer, one day beyond the minimum development time at this temperature, to maximize the proportion of emerging adults (>85% in all but four vials) while minimizing fight-related mortality. This timing ensured sufficient maturation across the majority of individuals prior to assessment. Tritonymphs were counted in the first counts for each generation.

These first adult counts were taken while transferring mites into new vials. Thereafter, heatwave populations were subjected to a temperature increase for 45 hours. The heatwave temperature initially started at 34°C (F_0_) and was increased by 1°C every two generations, reaching 38°C by the final generation. Second adult counts were conducted while simultaneously transferring mites to new vials for a 2-days mating/egg-laying period. Adult mites were sexed for both counts and fighter males distinguished from scramblers. Subsequently, the adults were killed, and the eggs were given four days to hatch. For the next generation, 50 larvae were typically transferred. In a few cases (3.17%), first adult counts slightly exceeded 50 (by 1 to 3 adults), and the number of larvae was changed to the number of adults in such cases to avoid exclusion. If fewer than 50 larvae were available, nymphs were included to reach a total of 50 individuals, or as close as possible. This protocol was repeated for each generation (Fig. 1).

A population was considered extinct if (a) there were no larvae or (b) there were either no adults or no females in the first count.

### Statistics

Survival data were initially analysed using generalized linear mixed effect models (GLMM) with binomial error structures. If diagnostics indicated issues with over-dispersion, GLMMs with a beta-binomial error distribution were used. The number of females (count data) was analysed using a linear mixed effects model (LMM). The glmmTMB package (Brooks et al., 2017) was used for fitting all the models, and diagnostic plots were generated using the DHARMa package (Brooks et al., 2024; Hartig, 2024). Statistical models included as fixed effects the pheromone and temperature regimes, and generation as a continuous variable, plus the two-way interactions between the fixed effects.

Population ID was entered as a random effect. Each model with a full set of explanatory variables and their interactions were reduced to minimal adequate models in which only significant explanatory variables were retained (Crawley, 2012). All analyses were performed in R version 4.4.1 (R Core Team, 2024).

To estimate the effectiveness of our pheromone treatment, the proportion of fighter morph in each count was analysed as a binary response variable, e.g. cbind(fighters, scamblers). For this we used the subset of populations for which all juveniles reached adulthood at the first count, as it was not possible to determine the morph of the tritonymphs. This limited our sample size to 68.3% of the total dataset. Count (first or second adult counts) was entered as an additional fixed effect.

For analysing the proportion of individuals surviving at each generation we used a binary response variable in the form of a vector containing the numbers of survived and dead individuals. The proportion survival inferred from two subsequent counts within a population, i.e. dead = the number initially present – number survived. It was analysed separately for juveniles to first adult counts (early survival) and first to second adult counts (late survival). F_0_ counts were not included in analyses of early survival, because this stage was not exposed to direct or cross-generation effects of heatwaves. For the sex and morph specific survival last two we used a subset of populations without tritonymph same as in the analysis of fighter proportions but with sex and morph as a fixed effect respectively instead of count. Sex specific survival was also the only model where we tested a three-way interaction (temperature*sex*generation) to see if there was a sex difference in survival over generations. We further analysed the proportion of females in the population only for the second adult counts to additionally see the indirect effects of fights on skewness in the sex ratio. For this, we used the proportion of females as a binary response variable. Furthermore, we analysed the absolute counts of females from the second count, as this number sets an upper limit on the productivity of the population for the next generation.

## Results

None of the populations in the stable temperatures went extinct. Of the heatwave treatments, no population went extinct in control treatment (high fighter prevalence), but three of the pheromone treatment (lower fighter prevalence) heatwave populations went extinct in the second, fourth and the sixth generation respectively. Two of the populations became extinct because there were no females in the first adult counts while the third one became extinct due to no adults (only one tritonymph) in the first adult counts. Additionally, for most of the populations we had 50 larvae to start the next generations except for 5.9% times in pheromone treatment with heatwaves.

### Fighter proportions

Pheromone exposure caused a reduction in the proportion of males developing into the fighter morph: the proportion of fighter males in the pheromone control was significantly higher than in the pheromone treatment, with ∼64% of males being fighters in pheromone control and ∼40% in pheromone treatment across the entire experiment (main effect of pheromone treatment: *χ*2 = 21.524, d.f. = 1, *p* < 0.001, Fig. 2), confirming the effectiveness of pheromone treatment. Additionally, there was a significant temperature*generation interaction indicating that the proportion of fighters increased over time in the heatwave treatments only (*χ*2 = 4.724, d.f. = 1, *p* = 0.030, Fig. 2).

**Fig. 2:**
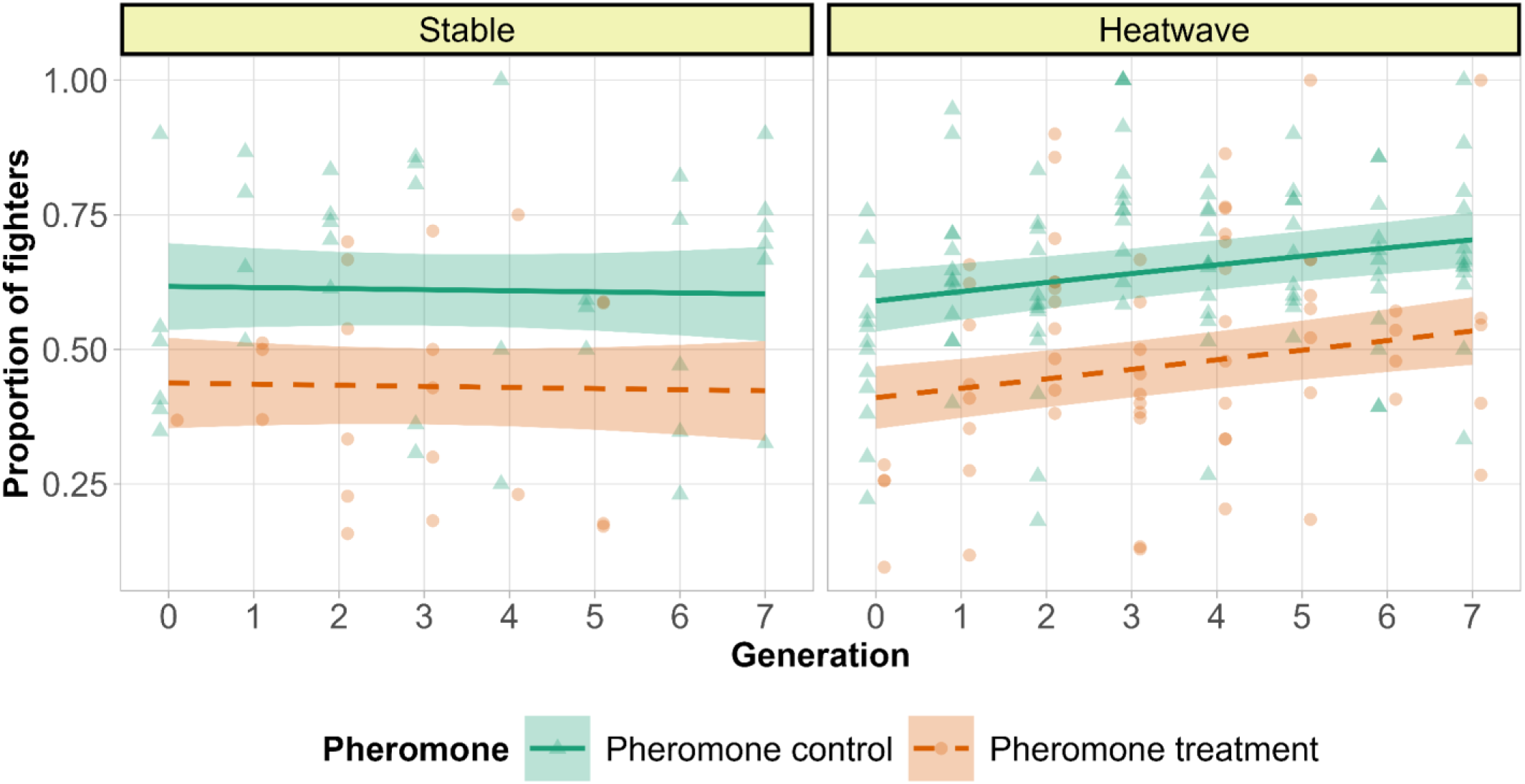
Fighter proportion in the control and heatwave treatments. Count was not retained in the final model, so for graphical representation of the model the average proportion of fighters from the two counts was presented. The lines are predicted by the model and points are raw data. They denote pheromone control (higher fighter prevalence) and orange dashes and circles denote pheromone treatment (lower fighter prevalence) with smooth areas as 95% confidence intervals.

### Survival

Late survival was higher in pheromone treatment than in pheromone control (*χ*2 = 10.115, d.f. = 1, *p* = 0.002) and the late survival was reduced by exposure to heatwaves (*χ*2 = 70.897, d.f. = 1, *p* = <0.001, Fig. 3b). Survival declined over generations in all treatments (*χ*2 =11.774, d.f. = 1, *p* = <0.001, late survival, Fig. 3). However, the decline in early survival for the pheromone treatment over generations was higher than for the pheromone control, as indicated by a significant pheromone * generation interaction, such that the initial higher survival in pheromone treatment populations largely disappeared towards the last generation (*χ*2 = 4.208, d.f. = 1, *p* = 0.040, Fig. 3a).

**Fig 3:**
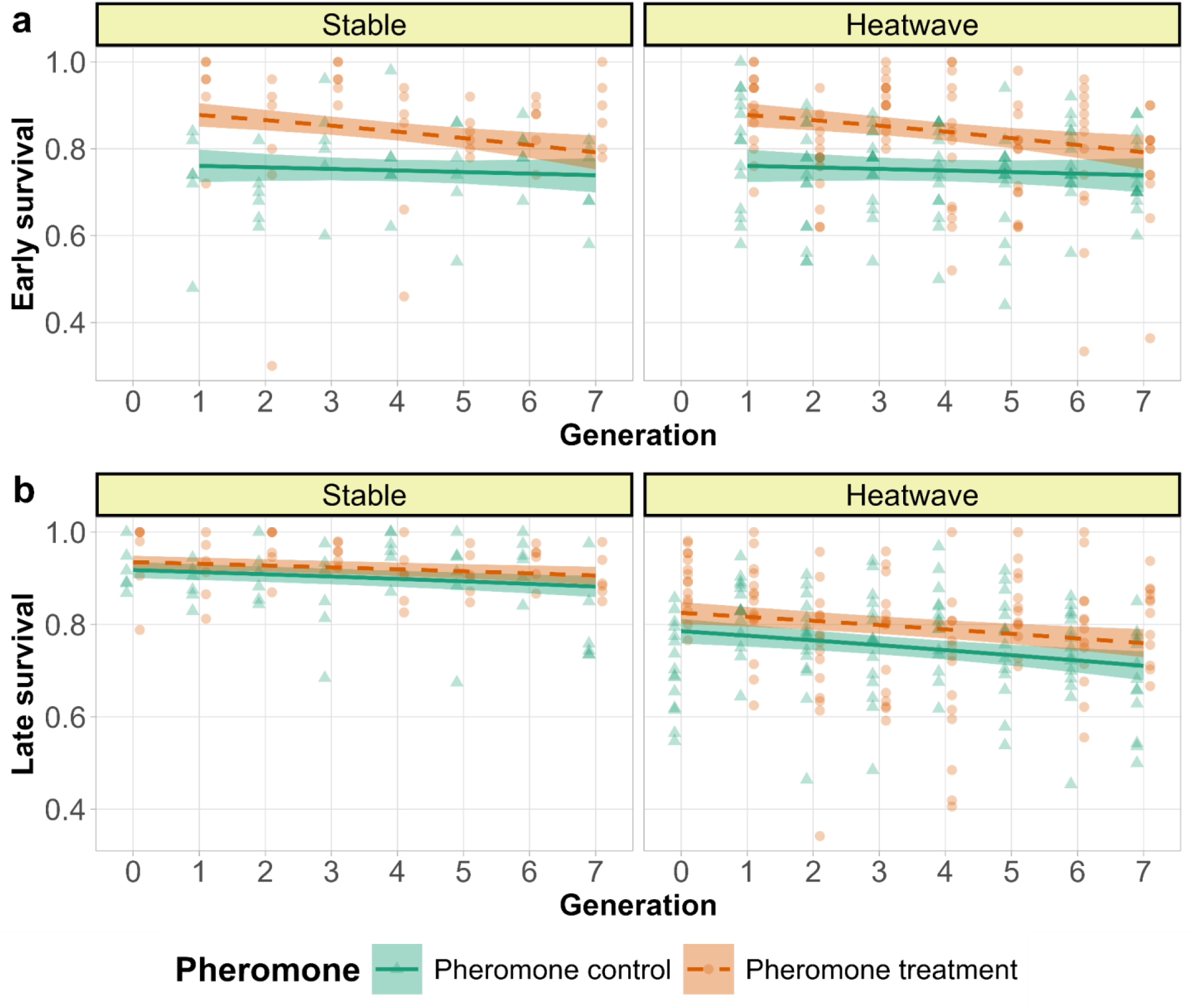
Proportion of survived individuals over generations in two temperature regimes indicated in the panels. a) Early survival b) Late survival over generations in temperature regimes in the panels. The lines are predicted by the model and points are raw data. The green lines and triangles denote pheromone control (higher fighter prevalence) and orange dashes and circles denote pheromone treatment (lower fighter prevalence). The smooth areas are 95% confidence intervals. Temperature regimes are indicated on the panels.

Females were more sensitive to heat than males, indicated by a significant two-way interaction between temperature and sex (*χ*2 = 23.957, d.f. = 1, *p* = <0.001, Fig. 4). Sex also interacted with pheromone treatment, with males, but not females, from pheromone treatment lines initially surviving better than in pheromone control lines (*χ*2 = 15.742, d.f. = 1, *p* = <0.001, Fig. 4). Furthermore, a significant sex*generation interaction indicates that female survival declined more than male survival over generations (*χ*2 = 6.187, d.f. = 1, *p* = 0.013, Fig. 4). Fighters in general survived better than scramblers (*χ*2 = 4.311, d.f. = 1, *p* = 0.038, Fig. 5).

**Fig. 4:**
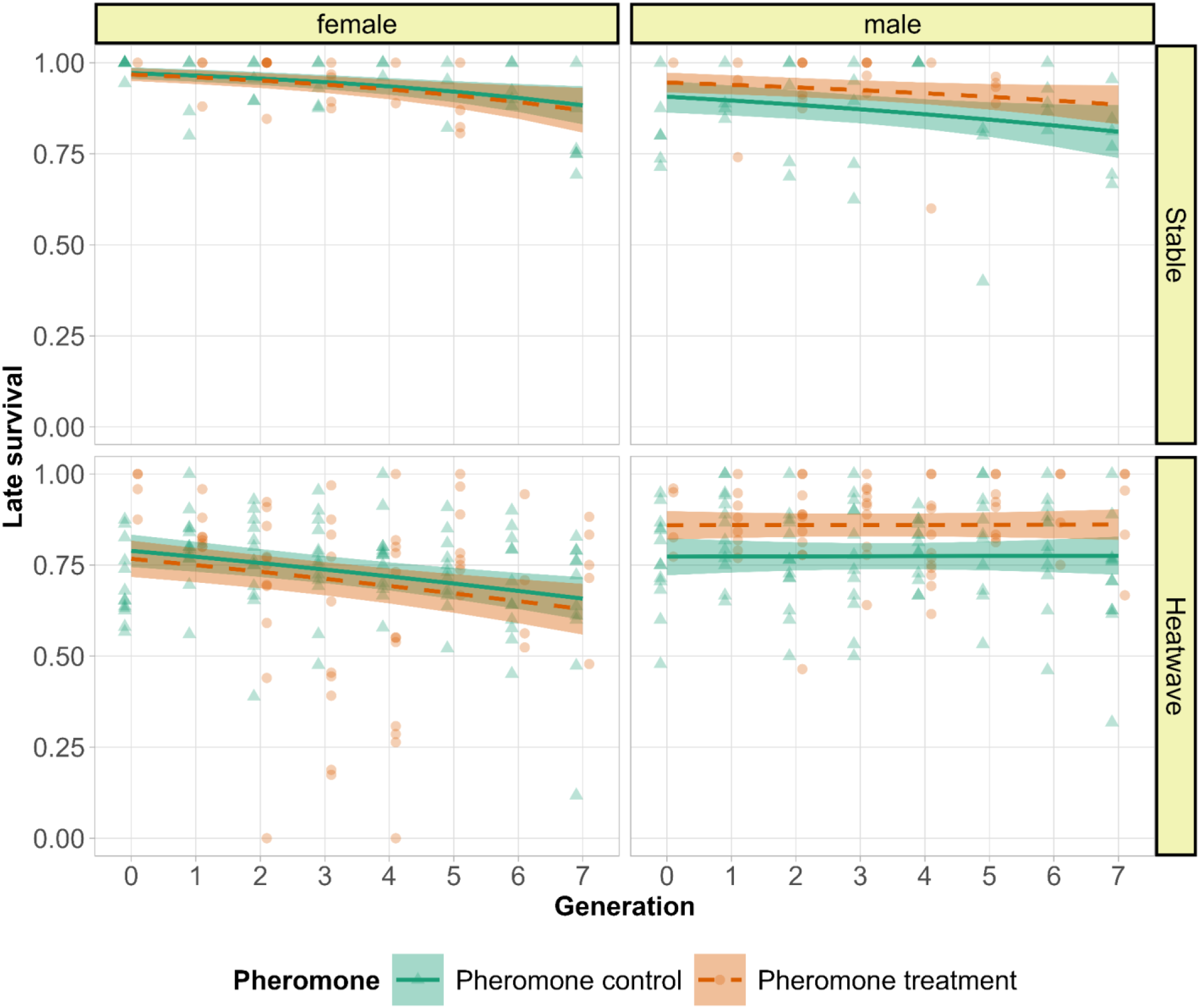
Sex-specific late survival. The lines are predicted by the model and points are raw data. The green lines and triangles denote pheromone control (higher fighter prevalence) and orange dashes and circles denote pheromone treatment (lower fighter prevalence) with smooth areas as 95% confidence intervals. The panels are sex in vertical and temperature regimes in horizontal.

**Fig. 5:**
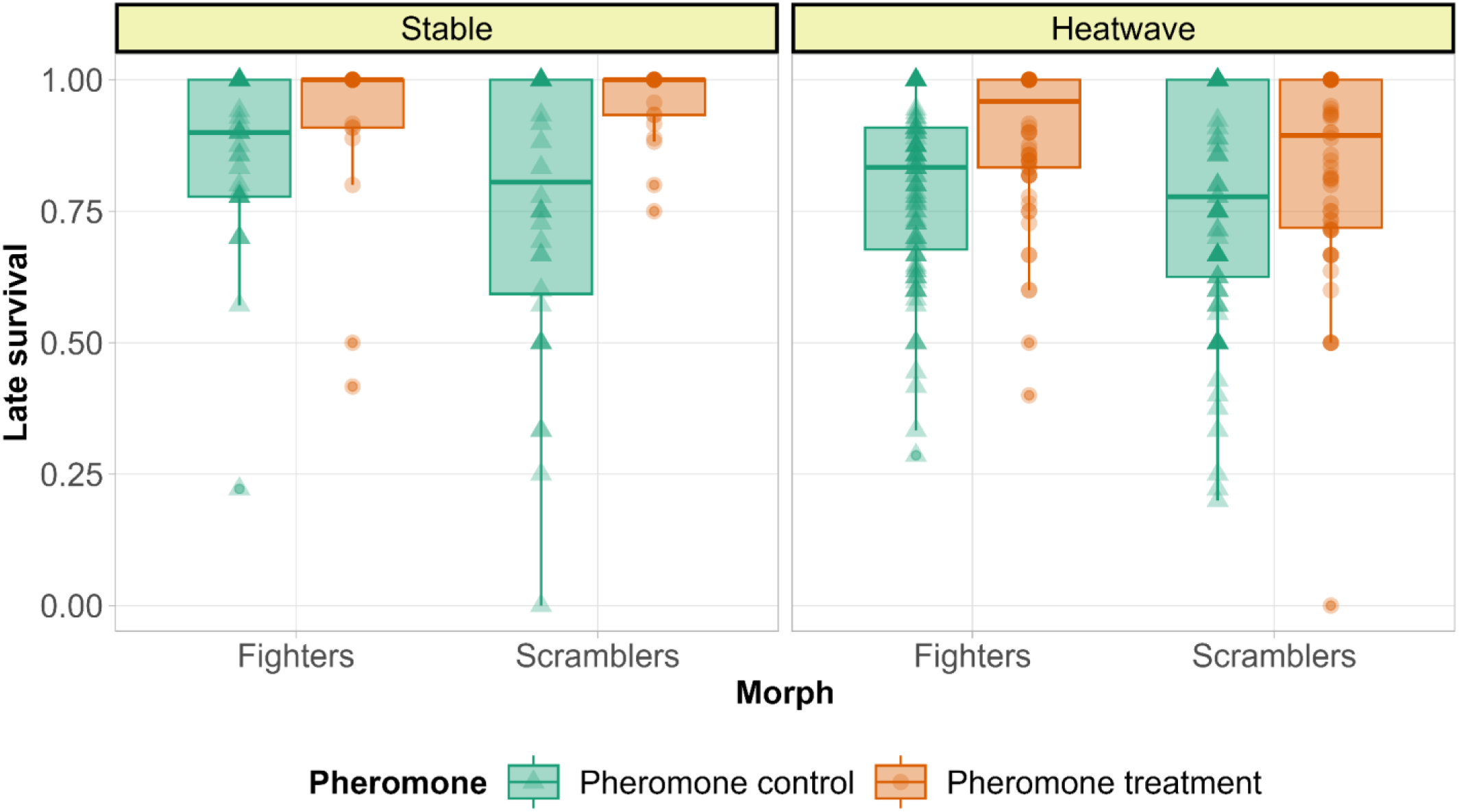
Morph specific late survival. The green boxes and triangles denote pheromone control (higher fighter prevalence) and orange boxes and circles denote pheromone treatment (lower fighter prevalence) and the panels are temperature regimes.

### Proportion and number of females at the onset of reproduction for larvae collection

Pheromone control populations were more female biased than the pheromone treatment (*χ*2 = 33.856, d.f. = 1, *p* = <0.001, Fig. 6). The number of females at second adult counts was negatively affected by heatwave regime (*F*= 33.678, *d.f.*= 1, *p*=<0.001, Fig. 6). Furthermore, there was a significant pheromone*generation interaction, with the decline in the number of females over generation being steeper in pheromone treatment than in pheromone control (*F*= 7.456, *d.f.*= 1, *p*=0.007, Fig. 6).

**Fig. 6:**
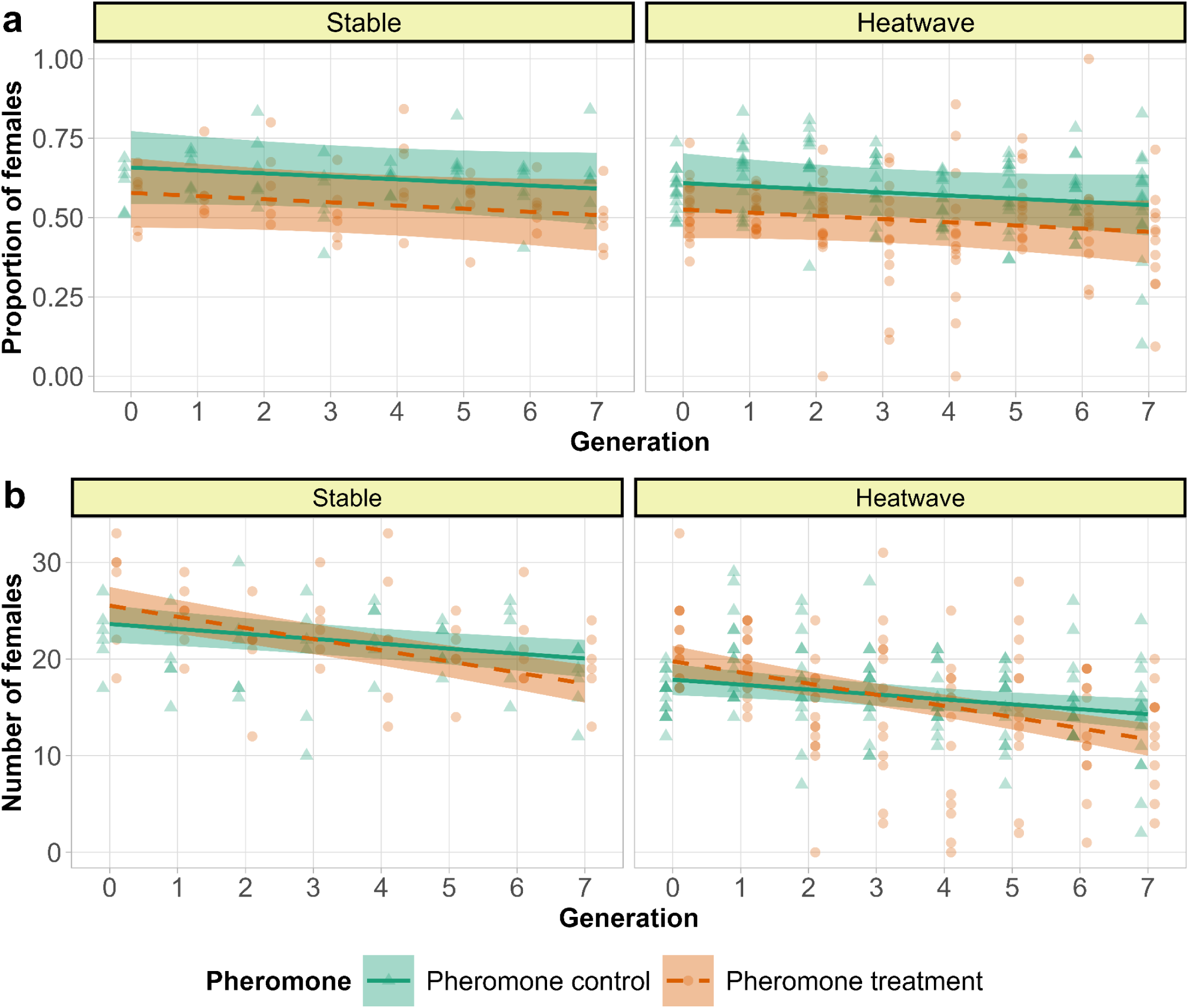
a) Proportion of females and b) Number of females in the second counts. The lines are predicted by the model and points are raw data. The green lines and triangles denote pheromone control (higher fighter prevalence) and orange dashes and circles denote pheromone treatment (lower fighter prevalence) with smooth areas as 95% confidence intervals. Temperature regimes are indicated on the panels.

## Discussion

The demographic consequences of strong sexual selection, particularly in small populations vulnerable to inbreeding and reduced adaptive capacity, are only beginning to be explored. In this study, we manipulated the intensity of sexual selection by suppressing the expression of armed, aggressive fighter morphs and examined the effects of heatwaves on survival. As predicted, both higher sexually selected trait expression and heatwaves reduced survival. These effects were sex-specific, with sexually selected trait expression primarily affecting males, while heatwaves had a stronger impact on females. We found no support, however, for our prediction that fighter males would experience disproportionately greater mortality under heatwaves because of the costs associated with weapon expression. We additionally found that survival declined over generations, independently of the direct effects of heatwaves. We discuss these findings and their broader implications in detail below.

Experimental heatwaves reduced late survival compared to controls. Sex-specific analysis showed that females were more sensitive to heatwaves than males, consistent with findings in *Tribolium castaneum* (Sales et al., 2021). However, in *R. robini*, and a more closely related species, no sex differences in survival during heatwaves were reported. A review by Edmands (2021) found no consistent sex-based differences in heat tolerance, but higher female resilience was more common.

Heatwaves reduced the number of reproductive females, a key factor in population dynamics. Independently, pheromone treatment affected male survival, with reduced survival in pheromone control populations where fighter morphs were more prevalent. This was accompanied by more female-biased sex ratios, suggesting elevated male mortality, likely due to lethal male fights (Radwan, 1993). However, contrary to our predictions, heat and pheromone treatments did not significantly interact in their effect on late survival. Contrary to the proposition that increased male mortality associated with expression of costly sexually selected weapons may be aggravated when conditions are poor (Bro-Jørgensen, 2014; Kokko & Brooks, 2003; Martínez-Ruiz & Knell, 2017), we found no differences between male morphs in their sensitivity to heat stress, consistent with the results in *R.robini* (Parrett et al., 2024). Furthermore, despite significantly female-biased sex ratios in higher fighter-prevalence populations, we noted no cases when there were no males at the mating/oviposition period. Additionally, we did not observe any extinctions in pheromone control heatwave populations, despite increased male mortality, even though we did note three extinctions in lower fighter-prevalence heatwave populations. Thus, despite associated reduced mortality, relaxation of sexual selection did not result in enhanced survival or decreased susceptibility to heatwaves.

Beyond the effects of heat and pheromone treatment, early and late survival declined over generations, as could be expected due to the negative impact of prolonged bottlenecks (Frankham, 2010; Spielman et al., 2004). Environmental challenges may exacerbate these effects (Reed et al., 2012), for which we found mixed support. While there was no significant temperature*generation interaction for early survival (Supplementary Table 2), we observed one for late survival, but only in a model accounting for sex (Supplementary Table 5). This may be because heatwave effects were sex-specific and ignoring this reduced statistical power by inflating within-population variance in survival.

Generational declines could possibly be mediated by sexual selection via reduced effective population sizes and increased range of inbreeding associated with increased male mortality.

However, the decline was not steeper in pheromone control populations, nor did it interact with heat. In contrast, early survival declined more steeply in pheromone treatment populations, eroding their initial survival advantage by the last generation (Fig. 3a). A similar interaction between pheromone treatment and generation was also observed for the number of females surviving till experimental egg-laying, which was initially lower in pheromone control treatment but with the rank reversed at the end of the experiment (Fig. 6). These results demonstrate that long-term positive effects of sexual selection on population dynamics in small populations can at least compensate for the negative effects of higher mortality it induces. The compensation is likely due to stronger selection against recessive deleterious alleles revealed by progressing inbreeding (Charlesworth & Willis, 2009). Indeed, enhanced purging of inbreeding load via sexual selection manipulated by enforcing monogamy have been demonstrated in studies in another acarid mite *R. robini* (Jarzebowska & Radwan, 2009) and in flour beetles *Tribolium castaneum* (Lumley et al., 2015). Our data also suggest that increasing the prevalence of a sexually selected weapon in populations may have a similar effect and corroborate the findings of fighter selected populations purging genetic load in *R. robini* (Parrett et al 2022).

Our results contrast with those obtained in another male-dimorphic acarid, *R. robini* (Łukasiewicz et al., 2023), which showed that fighter-selected populations exhibit poorer survival and greater extinction risk with rising temperatures. Here we observed only a few extinctions, but only in the heatwave regime with suppressed fighter expression. Additionally, irrespective of the heatwave regime, the initial survival advantage of these populations was decreasing over consecutive generations. One possible explanation for the contrasting conclusions is that the studies differed in the way heat stress was applied (heatwaves in the present study versus milder but permanent increase in Lukasiewicz et al. 2023). Another possible reason is the difference between the two species differ in the mode of male morph determination, being highly heritable in *R. robini* (Parrett et al., 2022; Radwan, 1995) but predominantly environmentally cued in *S. berlesei* (Michalczyk et al., 2018; Radwan, 1995). In *R. robini* morph determination maps to a large region of reduced recombination (Chmielewski et al. unpublished). Such regions can host sexually antagonistic loci (Jordan & Charlesworth, 2012), or accumulate deleterious variants (Carpentier et al., 2022; B. Charlesworth & Charlesworth, 2000), which can possibly affect how small populations respond to inbreeding and environmental stress. The effects of genetic architecture of sexually selected traits and the way the heat stress is applied deserves to be explored by future studies.

To conclude, our study shows that sexual selection can influence the survival of small populations. While high fighter prevalence increased male mortality and initially reduced the number of reproducing females, it also enhanced resilience to prolonged bottlenecks, likely through purging of inbreeding load. However, our results do not support the hypothesis that sexual selection increases vulnerability to environmental challenges. Contrary to theoretical models (Kokko & Brooks, 2003; Martínez-Ruiz & Knell, 2017), the negative effects of costly sexually selected traits were not exacerbated by environmental stress, nor did heatwaves interact with sexual selection or prolonged bottlenecks.We cannot, of course, rule out that increased male mortality from fights, combined with heatwave-driven female losses, would ultimately reduce long-term survival in small populations under strong sexual selection and demographic stochasticity (Martínez-Ruiz & Knell, 2017), although we only observed extinctions when fighter prevalence was low, indicating that this risk might be smaller than previously thought.

## Author contributions

J Radwan, N Porwal, JM Parrett, TC Cameron and RJ Knell conceived the ideas and designed methodology; N Porwal, A Szubert-Kruszyńska and N Pandey collected the data; N Porwal analysed the data with support from J Radwan, JM Parrett and RJ Knell; N Porwal and J Radwan led the writing of the manuscript with contributions from JM Parrett, TC Cameron and RJ Knell. All authors contributed critically to the drafts and gave final approval for publication.

## Acknowledgements

The project was funded by grant from NCN UMO-2020/39/B/NZ8/00152/4 to J Radwan, RJ Knell and TC Cameron.

