## Supplementary material for "Fighting Through the Heat: How Sexual Selection Influences Demography Under Recurrent Heatwaves"

**Table 1: Results from GLMM to predict fighter proportions without including the first and second adult counts where first adult counts had tritonymphs.**
*The factors and interactions retained in the final model are denoted in bold. $\chi^2$ values, d.f., and p-values are from likelihood ratio tests comparing models with and without the variable in question.*

| *Source of variation* | *$\chi^2$* | *d.f.* | *p-value* |
|---|---|---|---|
| *temperature* | - | - | - |
| ***pheromone*** | **21.524** | **1** | **<0.001** |
| *generation* | - | - | - |
| ***temperature * generation*** | **4.724** | **1** | **0.030** |
| *count* | 0.338 | 1 | 0.561 |
| *pheromone * count* | 0.035 | 1 | 0.852 |
| *temperature * pheromone* | 0.034 | 1 | 0.853 |
| *count * generation* | 0.018 | 1 | 0.894 |
| *pheromone * generation* | 0.008 | 1 | 0.930 |
| *temperature * count* | 0.003 | 1 | 0.956 |

**Table 2: Summary results from the model for fighter proportion**

| *Explanatory variable* | *est.* | *s.e.* | *z-value* | *p-value* |
|---|---|---|---|---|
| *Intercept* | 0.476 | 0.174 | 2.737 | **0.006** |
| *temperatureheatwave* | -0.111 | 0.189 | -0.589 | 0.556 |
| *pheromonetreatment* | -0.727 | 0.136 | -5.356 | **<0.001** |
| *generation* | -0.008 | 0.031 | -0.267 | 0.789 |
| *temperatureheatwave:generation* | 0.080 | 0.037 | 2.180 | **0.029** |

**Table 3: Results from fitting GLMMs to predict individual early and late survival.**
*The factors and interactions retained in the final model are denoted in bold. $\chi^2$ values, d.f., and p-values are from likelihood ratio tests comparing models with and without the variable in question.*

| *Source of variation* | *Early Survival* | | | *Late Survival* | | |
|---|---|---|---|---|---|---|
| | $\chi^2$ | d.f. | **p-value** | $\chi^2$ | d.f. | **p-value** |
| *temperature* | 1.947 | 1 | 0.163 | **70.897** | **1** | **<0.001** |
| *pheromone* | - | - | - | **10.115** | **1** | **0.002** |

| | | | | | | |
|---|---|---|---|---|---|---|
| *generation* | - | - | - | **11.774** | **1** | **<0.001** |
| ***pheromone * generation*** | **4.208** | **1** | **0.040** | 1.570 | 1 | 0.210 |
| *temperature * pheromone* | 3.433 | 1 | 0.064 | 0.490 | 1 | 0.484 |
| *temperature * generation* | 0.907 | 1 | 0.341 | 0.255 | 1 | 0.614 |

**Table 4: Model summary results from models for early and late survival**

| ***Explanatory variable*** | ***Early Survival*** | | | | ***Late Survival*** | | | |
|---|---|---|---|---|---|---|---|---|
| | est. | s.e. | z-value | p-value | est. | s.e. | z-value | p-value |
| *Intercept* | 1.176 | 0.127 | 9.264 | <0.001 | 2.413 | 0.117 | 22.554 | **<0.001** |
| *temperatureheat wave* | | | | | -1.113 | 0.100 | -11.106 | **<0.001** |
| *pheromonetreatment* | 0.904 | 0.200 | 4.516 | <0.001 | 0.253 | 0.077 | 3.301 | **<0.001** |
| *generation* | -0.019 | 0.027 | -0.716 | 0.474 | -0.057 | 0.017 | -3.453 | **<0.001** |
| *pheromonetreatment:generation* | -0.087 | 0.042 | -2.059 | 0.040 | | | | |

**Table 5: Results from fitting a GLMM to predict sex-specific late survival without including the counts where first adult counts had tritonymphs.**
*The factors and interactions retained in the final model are denoted in bold. $\chi^2$ values, d.f., and p-values are from likelihood ratio tests comparing models with and without the variable in question.*

| ***Source of variation*** | ***$\chi^2$*** | ***d.f.*** | ***p-value*** |
|---|---|---|---|
| *temperature* | - | - | - |
| *pheromone* | - | - | - |
| *generation* | - | - | - |
| *sex* | - | - | - |
| ***temperature * sex*** | **23.957** | **1** | **<0.001** |
| ***temperature * generation*** | **4.757** | **1** | **0.029** |
| ***pheromone * sex*** | **15.742** | **1** | **<0.001** |
| ***sex * generation*** | **6.187** | **1** | **0.013** |
| *pheromone * generation* | 1.377 | 1 | 0.241 |
| *temperature * pheromone* | 1.027 | 1 | 0.311 |
| *temperature * sex * generation* | 3.018 | 1 | 0.082 |

**Table 6: Summary results from the model for sex-specific survival after heatwave.**

| *Explanatory variable* | *est.* | *s.e.* | *z-value* | *p-value* |
|---|---|---|---|---|
| *Intercept* | 3.521 | 0.289 | 12.189 | **<0.001** |
| *temperatureheatwave* | -2.203 | 0.285 | -7.716 | **<0.001** |
| *pheromonetreatment* | -0.126 | 0.144 | 0.880 | 0.379 |
| *sexmale* | -1.248 | 0.271 | -4.602 | **<0.001** |
| *generation* | -0.214 | 0.055 | -3.873 | **<0.001** |
| *temperatureheatwave:sexmale* | 1.154 | 0.235 | 4.906 | **<0.001** |
| *temperatureheatwave:generation* | 0.119 | 0.055 | 2.174 | **<0.030** |
| *pheromonetreatment:sexmale* | 0.710 | 0.178 | 3.990 | **<0.001** |
| *sexmale:generation* | 0.097 | 0.040 | 2.491 | **<0.001** |

**Table 7: Results from fitting a GLMM to predict morph-specific late survival without including the counts where first adult counts had tritonymphs.**

*The factors and interactions retained in the final model are denoted in bold. $\chi^2$ values, d.f., and p-values are from likelihood ratio tests comparing models with and without the variable in question.*

| *Source of variation* | *$\chi^2$* | *d.f.* | *p-value* |
|---|---|---|---|
| ***temperature*** | **12.507** | **1** | **<0.001** |
| ***pheromone*** | **16.856** | **1** | **<0.001** |
| ***morph*** | **4.311** | **1** | **0.038** |
| *generation* | 4.532 | 1 | 0.270 |
| *temperature * pheromone* | 2.480 | 1 | 0.115 |
| *pheromone * generation* | 0.634 | 1 | 0.426 |
| *morph * generation* | 0.662 | 1 | 0.416 |
| *pheromone * morph* | 0.130 | 1 | 0.718 |
| *temperature * generation* | 0.121 | 1 | 0.728 |
| *temperature * morph* | 0.031 | 1 | 0.860 |

**Table 8: Model summary results from model for morph-specific survival**

| *Explanatory variable* | *est.* | *s.e.* | *z-value* | *p-value* |
|---|---|---|---|---|
| *Intercept* | 1.962 | 0.172 | 11.409 | **<0.001** |
| *temperatureheatwave* | -0.659 | 0.180 | -3.656 | **<0.001** |
| *pheromonetreatment* | 0.689 | 0.157 | 4.401 | **<0.001** |
| *morphscrambler* | -0.193 | 0.093 | -2.085 | **0.037** |

**Table 9: Results from fitting a GLMM and LMM to predict female proportion and female number respectively in the second counts.**
*The factors and interactions retained in the final model are denoted in bold. $\chi^2$, F values, d.f., and p-values are from likelihood ratio tests comparing models with and without the variable in question.*

| ***Source of variation*** | ***Female Proportion*** | | | ***Number of Females*** | | |
|---|---|---|---|---|---|---|
| | $\chi^2$ | d.f. | p | F value | d.f. | **p-value** |
| ***temperature*** | **14.153** | **1** | **<0.001** | **48.049** | **1** | **<0.001** |
| ***pheromone*** | **33.856** | **1** | **<0.001** | - | - | - |
| ***generation*** | **21.913** | **1** | **<0.001** | - | - | - |
| *temperature * generation* | 2.144 | 1 | 0.143 | 2.501 | 1 | 0.115 |
| *temperature * pheromone* | 1.373 | 1 | 0.241 | 3.133 | 1 | 0.085 |
| ***pheromone * generation*** | 1.180 | 1 | 0.274 | **7.456** | **1** | **0.007** |

**Table 10: Model summary results from model for proportion of females and number of females in the second counts**

| *Explanatory variable* | ***Proportion of Females*** | | | | ***Number of Females*** | | | |
|---|---|---|---|---|---|---|---|---|
| | est. | s.e. | z-value | p-value | est. | s.e. | t-value | p-value |
| *Intercept* | 0.653 | 0.058 | 11.092 | <0.001 | 23.616 | 0.979 | 24.119 | **<0.001** |
| *temperatureheatwave* | -0.211 | 0.052 | -4.094 | <0.001 | -5.757 | 0.831 | -6.932 | **<0.001** |
| *pheromonetreatment* | -0.339 | 0.048 | -7.006 | <0.001 | 1.9033 | 1.105 | 1.722 | **0.087** |
| *generation* | -0.040 | 0.009 | -4.674 | <0.001 | -0.510 | 0.164 | -3.117 | **0.002** |
| *pheromonetreatment: generation* | | | | | -0.644 | 0.236 | -2.731 | **0.007** |

**R script**

```
rm(list = ls(all = TRUE))
ls()

# packages ------------------------------------------------------------

library(readxl)
library(glmmTMB)
library(ggplot2)
library(bbmle)
library(pscl)
library(lmerTest)
library(DHARMa)
library(ggpubr)
library(ggh4x)
library(patchwork)
library(tidyr)
library(tidyverse)
library(dplyr)

setwd("~/Desktop/extinction_sancassania/final")

# function for overdispersion --------------------------------------------

overdisp_fun <- function(model) {
  rdf <- df.residual(model)
  rp <- residuals(model, type = "pearson")
  Pearson.chisq <- sum(rp ^ 2)
  prat <- Pearson.chisq / rdf
  pval <- pchisq(Pearson.chisq, df = rdf, lower.tail = FALSE)
```

```
  c(
    chisq = Pearson.chisq,
    ratio = prat,
    rdf = rdf,
    p = pval
  )
}

# Fighter proportions- adult survival ------------------------------------

rm(list = ls(all = TRUE))
ls()

san_morph = read_excel("SS_heat_demography.xlsx", sheet = "morph")
san_morph$line = as.factor(san_morph$line)
san_morph$Pheromone = as.factor(san_morph$Pheromone)
san_morph$Temperature <- as.factor(san_morph$Temperature)
san_morph$fig = as.numeric(san_morph$fig)
san_morph$scr = as.numeric(san_morph$scr)
san_morph$female = as.numeric(san_morph$female)
san_morph$gen = as.numeric(san_morph$gen)



san_morph_notrit = subset(san_morph, gen != 8)
san_morph_notrit = subset(san_morph_notrit, trito == 0)
san_morph_notrit$Temperature <- factor(san_morph_notrit$Temperature, levels = c("Stable",
"Heatwave"))



morph_wo_trit <- glmer(
```

```
  cbind(fig, scr) ~ (Temperature + Pheromone + count +
                     as.numeric(gen)) ^ 2  + (1 |line),
  data = san_morph_notrit,
  family = binomial,
  control = glmerControl(optimizer = 'bobyqa', optCtrl = list(maxfun = 100000))
)
drop1(morph_wo_trit, test = "Chi")

morph_wo_trit <- update(morph_wo_trit,~.-Temperature:Pheromone)
drop1(morph_wo_trit, test = "Chi")

morph_wo_trit <- update(morph_wo_trit,~.-Pheromone:as.numeric(gen))
drop1(morph_wo_trit, test = "Chi")

morph_wo_trit <- update(morph_wo_trit,~.-Temperature:count)
drop1(morph_wo_trit, test = "Chi")

morph_wo_trit <- update(morph_wo_trit,~.-count:as.numeric(gen))
drop1(morph_wo_trit, test = "Chi")

morph_wo_trit <- update(morph_wo_trit,~.-Pheromone:count)
drop1(morph_wo_trit, test = "Chi")

morph_wo_trit <- update(morph_wo_trit,~.-count)
drop1(morph_wo_trit, test = "Chi")

#check for overdispersion
overdisp_fun(morph_wo_trit)#ratio=1.44

#diagnostic plots
```

```
simulateResiduals(fittedModel = morph_wo_trit, plot = TRUE)

#accounting for overdispersion using betabinomial error distribution

morph_beta = glmmTMB(cbind(fig, scr) ~ (Temperature + Pheromone + count + as.numeric
(gen))^2 + (1|line), data = san_morph_notrit, family = betabinomial,
                  control=glmmTMBControl(optCtrl=list(iter.max=1000000)))
drop1(morph_beta,test="Chi")

morph_beta <- update(morph_beta,~.-Temperature:count)
drop1(morph_beta, test = "Chi")

morph_beta <- update(morph_beta,~.-Pheromone:as.numeric(gen))
drop1(morph_beta, test = "Chi")

morph_beta <- update(morph_beta,~.-count:as.numeric(gen))
drop1(morph_beta, test = "Chi")

morph_beta <- update(morph_beta,~.-Temperature:Pheromone)
drop1(morph_beta, test = "Chi")

morph_beta <- update(morph_beta,~.-Pheromone:count)
drop1(morph_beta, test = "Chi")

morph_beta = glmmTMB(cbind(fig, scr) ~ (Temperature + Pheromone + as.numeric(gen))^2
                  -Pheromone:as.numeric(gen)
                  -Temperature:Pheromone  + (1|line), data = san_morph_notrit, family =
betabinomial,
                  control=glmmTMBControl(optCtrl=list(iter.max=1000000)))
drop1(morph_beta,test="Chi")
```

```
#diagnostic plots
simulateResiduals(fittedModel = morph_beta, plot = TRUE)

summary(morph_beta)

#graph for fighter proportion

average_counts <- san_morph_notrit %>%
  filter(count %in% c("First adult count", "Second adult count")) %>%
  group_by(line, gen) %>%
  summarize(
    average_fig = mean(fig, na.rm = TRUE),
    average_scr = mean(scr, na.rm = TRUE),
    Pheromone = first(Pheromone),
    Temperature = first(Temperature),
    .groups = 'drop'
  )



average_counts$gen<- as.numeric(average_counts$gen)



# Update Pheromone labels to match your preferences
average_counts$fig_prop <- (average_counts$average_fig / (average_counts$average_fig +
average_counts$average_scr))

# write_xlsx(average_counts, path = "average fighter proportion.xlsx")

morph_dummy <- data.frame(
```

```
  gen = rep((0:7), times = 4),
  Temperature = rep(c("Stable", "Heatwave"), each = 16),
  #count = rep(c("First adult count", "Second adult count"), each = 16, times = ),
  Pheromone = rep(c("Control", "Treatment"), each = 8, times = 2)
)

predicted_morph <- predict(
  morph_beta,
  newdata = data.frame(morph_dummy, line = NA),
  se.fit = TRUE,
  type = "response"
)

# Create predicted values dataframe with the CI bounds
predicted_values_morph <- data.frame(morph_dummy,
                                     predict = predicted_morph$fit,
                                     se = predicted_morph$se.fit)

predicted_values_morph$lowerCI <- predicted_values_morph$predict - 1.96 *
predicted_values_morph$se
predicted_values_morph$upperCI <- predicted_values_morph$predict + 1.96 *
predicted_values_morph$se



predicted_values_morph$Temperature <- factor(predicted_values_morph$Temperature, levels =
c("Stable", "Heatwave"))

fig_2 <- ggplot(average_counts,
                aes(
                  x = as.numeric(gen),
                  y = fig_prop,
```

```
                  shape = Pheromone,  # Different shapes for points
                  linetype = Pheromone,  # Different line types for trends
                  colour = Pheromone,
                  fill = Pheromone
                )) +
geom_point(size = 2.2, alpha = 0.3, position = position_dodge(width =0.40)) +
scale_x_continuous(breaks = c(0, 1, 2, 3, 4, 5, 6, 7)) +

scale_x_continuous(breaks = c(0, 1, 2, 3, 4, 5, 6, 7)) +


scale_color_manual(values = c("#1b9e77", "#d95f02"),
                name = "Pheromone",
                labels = c("Pheromone control", "Pheromone treatment")) +
scale_fill_manual(values = c("#1b9e77", "#d95f02"),
               name = "Pheromone",
               labels = c("Pheromone control", "Pheromone treatment")) +
scale_shape_manual(values = c(17, 16),
                name = "Pheromone",
                labels = c("Pheromone control", "Pheromone treatment")) +
scale_linetype_manual(values = c("solid", "dashed"),
                  name = "Pheromone",
                  labels = c("Pheromone control", "Pheromone treatment")) +

# Add ribbon for CI shading
geom_ribbon(data = predicted_values_morph,
         aes(x = gen, ymin = lowerCI, ymax = upperCI, fill = Pheromone),
         alpha = 0.3, inherit.aes = FALSE) +

# Add the line for predicted values
geom_line(data = predicted_values_morph,
```

```
          aes(x = gen, y = predict, colour = Pheromone, linetype = Pheromone), linewidth = 1) +

  facet_grid(~Temperature) +
  labs(x = "Generation", y = "Proportion of fighters") +
  theme_light() +
  theme(panel.grid.minor = element_blank(),
        legend.key.size = unit(1, 'cm'),
        legend.key.height = unit(1, 'cm'),
        legend.key.width = unit(1, 'cm'),
        legend.title = element_text(size=14, face = "bold"),
        legend.text = element_text(size=14),
        legend.position = "bottom",
        axis.text = element_text(size = 14),
        axis.title = element_text(size = 14, face = "bold"),
        strip.text.x = element_text(size = 14, color = "black"),
        strip.text.y = element_text(size = 14, color = "black"),
        strip.background = element_rect(
          color = "black",
          fill = "#F2F4B5",
          size = 1.5,
          linetype = "solid"
        )
  )

# Display the plot
fig_2


ggsave(
  filename = "fig_2.pdf",
```

```
  plot = fig_2,
  dpi = 900
)
# Population level individual survival ------------------------------------

san_ext <- read_excel("SS_heat_demography.xlsx", sheet = "survival")

san_ext = subset(san_ext, generation != 8)

san_ext$line = as.factor(san_ext$line)
san_ext$morph = as.factor(san_ext$Pheromone)
san_ext$juvenile = as.numeric(san_ext$juvenile)
san_ext$ad_total_I = as.numeric(san_ext$ad_total_I)
san_ext$ad_total_II = as.numeric(san_ext$ad_total_II)

san_ext$Temperature <- factor(san_ext$Temperature, levels = c("Stable", "Heatwave"))

# Survival from juvenile to first adult count -----------------------------
san_juv_sur <- san_ext %>%
  mutate(across(c(ad_total_I, juvenile), ~ ifelse(generation == 0, NA, .)))

#san_juv_sur<- subset(san_juv, generation != 0)

juv_sur = glmer(
  cbind(ad_total_I, juvenile - ad_total_I) ~ (Temperature + Pheromone +
  as.numeric(generation)) ^ 2  + (1 | line),
  data = san_juv_sur,
  family = binomial,
  control = glmerControl(optimizer = 'bobyqa', optCtrl = list(maxfun = 100000))
)
```

```
drop1(juv_sur, test = "Chi")

juv_sur = glmer(cbind(ad_total_I, juvenile - ad_total_I) ~ (Temperature + Pheromone +
  as.numeric(generation)) ^ 2 - Temperature:Pheromone + (1 | line),
  data = san_juv_sur,
  family = binomial,
  control = glmerControl(optimizer = 'bobyqa', optCtrl = list(maxfun = 100000))
)
drop1(juv_sur, test = "Chi")

#diagnostic plots
simulateResiduals(fittedModel = juv_sur, plot = TRUE)

#checking for overdispersion
overdisp_fun(juv_sur) # ratio= 3.57

#accounting for over dispersion using beta-binomial error distribution

juv_sur_beta = glmmTMB(
  cbind(ad_total_I, juvenile - ad_total_I) ~ (Temperature + Pheromone +
  as.numeric(generation))^2 + (1 | line),
  data = san_juv_sur,
  family = betabinomial,
  control = glmmTMBControl(optCtrl = list(iter.max =
                                         1000000)))
drop1(juv_sur_beta, test = "Chi")

juv_sur_beta = glmmTMB(
  cbind(ad_total_I, juvenile - ad_total_I) ~ (Temperature + Pheromone +
```

```
                                    as.numeric(generation)) ^2 -Temperature:as.numeric(generation)+
(1 | line),
  data = san_juv_sur,
  family = betabinomial,
  control = glmmTMBControl(optCtrl = list(iter.max =
                                    1000000))
)
drop1(juv_sur_beta, test = "Chi")

juv_sur_beta = glmmTMB(
  cbind(ad_total_I, juvenile - ad_total_I) ~ (Temperature + Pheromone +
                                    as.numeric(generation)) ^2 -Temperature:as.numeric(generation) -
Temperature:Pheromone  + (1 | line),
  data = san_juv_sur,
  family = betabinomial,
  control = glmmTMBControl(optCtrl = list(iter.max =
                                    1000000)) # for graph
)
drop1(juv_sur_beta, test = "Chi")

juv_sur_beta = glmmTMB(
  cbind(ad_total_I, juvenile - ad_total_I) ~ (Pheromone +
                                    as.numeric(generation)) ^2  + (1 | line),
  data = san_juv_sur,
  family = betabinomial,
  control = glmmTMBControl(optCtrl = list(iter.max =
                                    1000000))
)
drop1(juv_sur_beta, test = "Chi")
```

```
#diagnostic plots
simulateResiduals(fittedModel = juv_sur_beta, plot = TRUE)

summary(juv_sur_beta) #best

#plotting the predictions from juvenile survival/ pre heat stress survival from juv_sur_beta
#create a data frame to store the predictions
juv_dummy <- data.frame(
  generation = rep((0:7), times = 4),
  Temperature = rep(c("Stable", "Heatwave"), each = 16),
  Pheromone = rep(c("Control", "Treatment"), each = 8, times = 2)
)

# Predict juvenile survival
predicted_juv <- predict(
  juv_sur_beta,
  newdata = data.frame(juv_dummy, line = NA),
  se.fit = TRUE,
  type = "response"
)

# Store predicted values and CIs
predicted_values_juv <- data.frame(juv_dummy, predict = predicted_juv$fit, se =
predicted_juv$se.fit)
predicted_values_juv$lowerCI <- predicted_values_juv$predict - 1.96 * predicted_values_juv$se
predicted_values_juv$upperCI <- predicted_values_juv$predict + 1.96 *
predicted_values_juv$se

predicted_values_juv$Temperature <- factor(predicted_values_juv$Temperature, levels =
c("Stable", "Heatwave"))
```

```
# Calculate the proportion of juveniles survived (raw data)
san_juv_sur$juv_prop = san_juv_sur$ad_total_I / san_juv_sur$juvenile

fig_3a <- ggplot(san_juv_sur,
                 aes(
                   x = as.numeric(generation),
                   y = juv_prop,
                   shape = Pheromone,  # Set shape for points
                   linetype = Pheromone,  # Set line type for lines
                   colour = Pheromone,
                   fill = Pheromone
                 )) +
  geom_point(size = 2.2, alpha = 0.3, position = position_dodge(width =0.40)) +
  scale_x_continuous(breaks = c(0, 1, 2, 3, 4, 5, 6, 7)) +

  scale_color_manual(values = c("#1b9e77", "#d95f02"),
                     name = "Pheromone",
                     labels = c("Pheromone control", "Pheromone treatment")) +
  scale_fill_manual(values = c("#1b9e77", "#d95f02"),
                    name = "Pheromone",
                    labels = c("Pheromone control", "Pheromone treatment")) +
  scale_shape_manual(values = c(17, 16),
                     name = "Pheromone",
                     labels = c("Pheromone control", "Pheromone treatment")) +
  scale_linetype_manual(values = c("solid", "dashed"),
                        name = "Pheromone",
                        labels = c("Pheromone control", "Pheromone treatment")) +

  # Add ribbon for CI shading (excluding generation == 0)
  geom_ribbon(data = predicted_values_juv %>% filter(generation > 0),
```

```
            aes(x = generation, ymin = lowerCI, ymax = upperCI, fill = Pheromone),
            alpha = 0.4, inherit.aes = FALSE) +

  # Add the line for predicted values (excluding generation == 0)
  geom_line(data = predicted_values_juv %>% filter(generation > 0),
           aes(x = generation, y = predict, colour = Pheromone, linetype = Pheromone), linewidth =
1) +

  facet_wrap(~ Temperature) +
  labs(x = "Generation", y = "Early survival") +
  theme_light() +
  theme(panel.grid.minor = element_blank()) +
  theme(
    axis.text = element_text(size = 16),
    axis.title = element_text(size = 16, face = "bold"),
    #axis.title.x = element_blank(), axis.text.x = element_blank(),
    strip.text.x = element_text(size = 16, color = "black"),
    strip.text.y = element_text(size = 16, color = "black"),
    strip.background = element_rect(
      color = "black", fill = "#F2F4B5", size = 1.5, linetype = "solid"
    ),
    legend.text = element_text(size = 15),
    legend.title = element_text(size = 18, face = "bold"),
    legend.key.height = unit(1.0, "cm"),
    legend.key.width = unit(1.0, "cm")
  )

# Display the plot
fig_3a
```

```
# First adult count to second adult count survival ------------------------

adult_sur = glmer(
  cbind(ad_total_II, ad_total_I - ad_total_II) ~ (Temperature + Pheromone +
  as.numeric(generation)) ^ 2   + (1 | line),
  data = san_ext,
  family = binomial,
  control = glmerControl(optimizer = 'bobyqa', optCtrl = list(maxfun = 100000))
)
drop1(adult_sur, test = "Chi")

adult_sur <- update(adult_sur,~.-Temperature:as.numeric(generation))
drop1(adult_sur, test = "Chi")

adult_sur <- update(adult_sur,~.-Temperature:Pheromone)
drop1(adult_sur, test = "Chi")

adult_sur <- update(adult_sur,~.-Pheromone:as.numeric(generation))
drop1(adult_sur, test = "Chi")

#check for overdispersion
overdisp_fun(adult_sur) # ratio= 2.4

#diagnostic plots
simulateResiduals(fittedModel = adult_sur, plot = TRUE)

#accounting for over dispersion using beta binomial error distribution

adult_sur_beta = glmmTMB(
```

```
  cbind(ad_total_II, ad_total_I - ad_total_II) ~ (Temperature + Pheromone +
  as.numeric(generation)) ^ 2 + (1 |line),
  data = san_ext,
  family = betabinomial,
  control = glmmTMBControl(optCtrl = list(iter.max =
                                        1000000))
)

drop1(adult_sur_beta, test = "Chi")

adult_sur_beta <- update(adult_sur_beta,~.-Temperature:as.numeric(generation))
drop1(adult_sur_beta, test = "Chi")

adult_sur_beta <- update(adult_sur_beta,~.-Temperature:Pheromone)
drop1(adult_sur_beta, test = "Chi")

adult_sur_beta = glmmTMB(
  cbind(ad_total_II, ad_total_I - ad_total_II) ~ (Temperature + Pheromone +
                                        as.numeric(generation)) ^ 2
  - Temperature:as.numeric(generation) -Temperature:Pheromone
  -Pheromone:as.numeric(generation)+ (1 |line),
  data = san_ext,
  family = betabinomial,
  control = glmmTMBControl(optCtrl = list(iter.max =
                                        1000000))
)
drop1(adult_sur_beta, test = "Chi")

#diagnostic plots
simulateResiduals(fittedModel = adult_sur_beta, plot = TRUE)
```

```
summary(adult_sur_beta)

#adults survived proportion
san_ext$adult_prop = san_ext$ad_total_II / san_ext$ad_total_I

# Prepare dummy data for predictions
adult_dummy <- data.frame(
  generation = rep((0:7), times = 4),
  Temperature = rep(c("Stable", "Heatwave"), each = 16),
  Pheromone = rep(c("Control", "Treatment"), each = 8, times = 2)
)

# Make predictions from the model (adult_sur_beta)
predicted_adult <- predict(
  adult_sur_beta,
  data.frame(adult_dummy, line = NA),
  se.fit = TRUE,
  type = "response"
)

# Store predicted values and their standard errors
predicted_values_adult <- data.frame(adult_dummy, predict = predicted_adult$fit, se =
predicted_adult$se.fit)
predicted_values_adult$lowerCI <- predicted_values_adult$predict - 1.96 *
predicted_values_adult$se
predicted_values_adult$upperCI <- predicted_values_adult$predict + 1.96 *
predicted_values_adult$se

san_ext$Temperature <- factor(san_ext$Temperature, levels = c("Stable", "Heatwave"))
```

```
predicted_values_adult$Temperature <- factor(predicted_values_adult$Temperature, levels =
c("Stable", "Heatwave"))

# Plot adult survival with predicted values and confidence intervals
fig_3b <- ggplot(san_ext,
                 aes(
                   x = as.numeric(generation),
                   y = adult_prop,
                   shape = Pheromone,  # Set shape for points
                   linetype = Pheromone,  # Set line type for lines
                   colour = Pheromone,
                   fill = Pheromone
                 )) +
  geom_point(size = 2.2, alpha = 0.3, position = position_dodge(width =0.40)) +
  scale_x_continuous(breaks = c(0, 1, 2, 3, 4, 5, 6, 7)) +

  scale_color_manual(values = c("#1b9e77", "#d95f02"),
                     name = "Pheromone",
                     labels = c("Pheromone control", "Pheromone treatment")) +
  scale_fill_manual(values = c("#1b9e77", "#d95f02"),
                    name = "Pheromone",
                    labels = c("Pheromone control", "Pheromone treatment")) +
  scale_shape_manual(values = c(17, 16),
                     name = "Pheromone",
                     labels = c("Pheromone control", "Pheromone treatment")) +
  scale_linetype_manual(values = c("solid", "dashed"),
                        name = "Pheromone",
                        labels = c("Pheromone control", "Pheromone treatment")) +

  # Add ribbon for CI shading
```

```
  geom_ribbon(data = predicted_values_adult,
              aes(x = generation, ymin = lowerCI, ymax = upperCI, fill = Pheromone),
              alpha = 0.4, inherit.aes = FALSE) +

  # Add the line for predicted values
  geom_line(data = predicted_values_adult,
            aes(x = generation, y = predict, colour = Pheromone, linetype = Pheromone), linewidth =
1) +

  facet_wrap(~ Temperature) +
  labs(x = "Generation", y = "Late survival") +
  theme_light() +
  theme(
    panel.grid.minor = element_blank(),
    axis.text = element_text(size = 16),
    axis.title = element_text(size = 16, face = "bold"),
    #axis.title.x = element_blank(),
    #axis.text.x = element_blank(),
    strip.text.x = element_text(size = 16, color = "black"),
    strip.text.y = element_text(size = 16, color = "black"),
    strip.background = element_rect(
      color = "black", fill = "#F2F4B5", size = 1.5, linetype = "solid"
    ),
    legend.text = element_text(size = 15),
    legend.title = element_text(size = 18, face = "bold"),
    legend.key.height = unit(1.0, "cm"),
    legend.key.width = unit(1.0, "cm")
  )

# Display the plot
```

```
fig_3b

fig_3<-ggarrange(
   fig_3a,
   fig_3b,
  nrow = 2,
  labels = c('a', 'b'),
  common.legend = TRUE,
  legend = "bottom",
  font.label = list(size = 20, face = "bold")
)

fig_3
ggsave(
  filename = "fig_3.pdf",
  plot = fig_3,
  dpi = 800,
)


# Sex specific adult survival --------------------------------------------

rm(list = ls(all = TRUE))
ls()

san_sex = read_excel("SS_heat_demography.xlsx", sheet = "sex")
san_sex$line = as.factor(san_sex$line)
san_sex$Pheromone = as.factor(san_sex$Pheromone)
san_sex$Temperature = as.factor(san_sex$Temperature)
san_sex$sex = as.factor(san_sex$sex)
```

```
san_sex$ad_total_I = as.numeric(san_sex$ad_total_I)
san_sex$ad_total_II = as.numeric(san_sex$ad_total_II)

san_sex_notrit = subset(san_sex, trito == 0)
san_sex_notrit = subset(san_sex, generation != 8)

san_sex_notrit$Temperature <- factor(san_sex_notrit$Temperature, levels = c("Stable",
"Heatwave"))

sex_sur = glmer(
  cbind(ad_total_II, ad_total_I - ad_total_II) ~ (Temperature + Pheromone + sex +
  as.numeric(generation))^2 + Temperature:sex:as.numeric(generation) + (1 |line),
  data = san_sex_notrit,
  family = binomial,
  control = glmerControl(optimizer = 'bobyqa', optCtrl = list(maxfun = 100000))
)
drop1(sex_sur, test = "Chi")

overdisp_fun(sex_sur)#ratio=1.9

#diagnostic plots
simulateResiduals(fittedModel = sex_sur, plot = TRUE)

#dealing with overdispersion
sex_sur_beta = glmmTMB(
  cbind(ad_total_II, ad_total_I - ad_total_II) ~ (Temperature + Pheromone + sex +
  as.numeric(generation)) ^2+ Temperature:sex:as.numeric(generation) + (1 | line),
  data = san_sex_notrit,
  family = betabinomial,
  control = glmmTMBControl(optCtrl = list(iter.max =
```

```
                                        10000)))
drop1(sex_sur_beta, test = "Chi")

sex_sur_beta <- update(sex_sur_beta,~.-Temperature:sex:as.numeric(generation))
drop1(sex_sur_beta, test = "Chi")

sex_sur_beta <- update(sex_sur_beta,~.-Temperature:Pheromone)
drop1(sex_sur_beta, test = "Chi")

sex_sur_beta = glmmTMB(
  cbind(ad_total_II, ad_total_I - ad_total_II) ~ (Temperature + Pheromone + sex +
                              as.numeric(generation)) ^2
  -Temperature:Pheromone -Pheromone:as.numeric(generation) + (1 |line),
  data = san_sex_notrit,
  family = betabinomial,
  control = glmmTMBControl(optCtrl = list(iter.max =
                                        10000))
)
drop1(sex_sur_beta, test = "Chi")

#diagnostic plots
simulateResiduals(fittedModel = sex_sur_beta, plot = TRUE)

san_sex_notrit$sex_prop_sur = san_sex_notrit$ad_total_II / san_sex_notrit$ad_total_I

summary(sex_sur_beta)

sex_dummy <- data.frame(
  generation = rep((0:7), times = 8),
  Temperature = rep(c("Stable", "Heatwave"), each = 32),
```

```
  sex = rep(c("male", "female"), each = 16, times = 2),
  Pheromone = rep(c("Control", "Treatment"), each = 8, times = 4)
)

predicted_sex <- predict(
  sex_sur_beta,
  newdata = data.frame(sex_dummy, line = NA),
  se.fit = TRUE,
  type = "response"
)

# Create predicted values dataframe with the CI bounds
predicted_values_sex <- data.frame(sex_dummy, predict = predicted_sex$fit, se =
predicted_sex$se.fit)
predicted_values_sex$lowerCI <- predicted_values_sex$predict - 1.96 * predicted_values_sex$se
predicted_values_sex$upperCI <- predicted_values_sex$predict + 1.96 *
predicted_values_sex$se

predicted_values_sex$Temperature <- factor(predicted_values_sex$Temperature, levels =
c("Stable", "Heatwave"))

# Plot sex specific survival with predicted values and confidence intervals
fig_4 <- ggplot(san_sex_notrit,
          aes(
            x = as.numeric(generation),
            y = sex_prop_sur,
            shape = Pheromone,  # Different shapes for points
            linetype = Pheromone,  # Different line types for trends
            colour = Pheromone,
            fill = Pheromone
          )) +
```

```
geom_point(size = 2.2, alpha = 0.3, position = position_dodge(width =0.40)) +
scale_x_continuous(breaks = c(0, 1, 2, 3, 4, 5, 6, 7)) +

scale_color_manual(values = c("#1b9e77", "#d95f02"),
                   name = "Pheromone",
                   labels = c("Pheromone control", "Pheromone treatment")) +
scale_fill_manual(values = c("#1b9e77", "#d95f02"),
                  name = "Pheromone",
                  labels = c("Pheromone control", "Pheromone treatment")) +
scale_shape_manual(values = c(17, 16),
                   name = "Pheromone",
                   labels = c("Pheromone control", "Pheromone treatment")) +
scale_linetype_manual(values = c("solid", "dashed"),
                      name = "Pheromone",
                      labels = c("Pheromone control", "Pheromone treatment")) +

# Add ribbon for CI shading
geom_ribbon(data = predicted_values_sex,
            aes(x = generation, ymin = lowerCI, ymax = upperCI, fill = Pheromone),
            alpha = 0.4, inherit.aes = FALSE) +

# Add the line for predicted values
geom_line(data = predicted_values_sex,
          aes(x = generation, y = predict, colour = Pheromone, linetype = Pheromone), linewidth =
1) +

facet_grid(Temperature ~ sex) +
labs(x = "Generation", y = " Late survival") +
theme_light() +
theme(panel.grid.minor = element_blank(),
```

```
        legend.key.size = unit(1, 'cm'),
        legend.key.height = unit(1, 'cm'),
        legend.key.width = unit(1, 'cm'),
        legend.title = element_text(size=14, face = "bold"),
        legend.text = element_text(size=14),
        legend.position = "bottom",
        axis.text = element_text(size = 14),
        axis.title = element_text(size = 14, face = "bold"),
        strip.text.x = element_text(size = 14, color = "black"),
        strip.text.y = element_text(size = 14, color = "black"),
        strip.background = element_rect(
          color = "black",
          fill = "#F2F4B5",
          size = 1.5,
          linetype = "solid"
        )
  )

# Display the graph
fig_4

ggsave(
  filename = "fig_4.pdf",
  plot = fig_4,
  dpi = 900
)


 # morph specific survival -----------------------------------
```

```
rm(list = ls(all = TRUE))
ls()

san_morph = read_excel("SS_heat_demography.xlsx", sheet = "morph")
san_morph$line = as.factor(san_morph$line)
san_morph$Pheromone = as.factor(san_morph$Pheromone)
san_morph$Temperature <- as.factor(san_morph$Temperature)
san_morph$fig = as.numeric(san_morph$fig)
san_morph$scr = as.numeric(san_morph$scr)
san_morph$gen = as.numeric(san_morph$gen)



san_morph_notrit = subset(san_morph, gen != 8)
san_morph_notrit = subset(san_morph_notrit, trito == 0)

san_morph_notrit$Temperature <- factor(san_morph_notrit$Temperature, levels = c("Stable",
"Heatwave"))

morph_sur <- san_morph_notrit %>%
  select(count, fig, scr, line, gen, Pheromone, Temperature) %>%
  pivot_longer(cols = c(scr, fig), names_to = "morph_type") %>%
  pivot_wider(
    id_cols = c(line, morph_type, gen, Pheromone, Temperature),
    names_from = count,
    values_from = value
  )



morph_sur_mod = glmer(
```

```
  cbind(`Second adult count`, (`First adult count` - `Second adult count`)) ~ (Temperature + Pheromone + morph_type +
                                as.numeric(gen))^2 + (1 |line),
  data = morph_sur,
  family = binomial,
  control = glmerControl(optimizer = 'bobyqa', optCtrl = list(maxfun = 100000))
)
drop1(morph_sur_mod, test = "Chi")

morph_sur_mod <- update(morph_sur_mod,~.-Temperature:morph_type)
drop1(morph_sur_mod, test = "Chi")

morph_sur_mod <- update(morph_sur_mod,~.-Temperature:as.numeric(gen))
drop1(morph_sur_mod, test = "Chi")

morph_sur_mod <- update(morph_sur_mod,~.-Pheromone:morph_type)
drop1(morph_sur_mod, test = "Chi")

morph_sur_mod <- update(morph_sur_mod,~.-morph_type:as.numeric(gen))
drop1(morph_sur_mod, test = "Chi")

morph_sur_mod <- update(morph_sur_mod,~.-Pheromone:as.numeric(gen))
drop1(morph_sur_mod, test = "Chi")

morph_sur_mod <- update(morph_sur_mod,~.-Temperature:Pheromone)
drop1(morph_sur_mod, test = "Chi")

morph_sur_mod = glmer(
  cbind(`Second adult count`, (`First adult count` - `Second adult count`)) ~ Temperature + Pheromone + morph_type + (1 |line),
```

```
  data = morph_sur,
  family = binomial,
  control = glmerControl(optimizer = 'bobyqa', optCtrl = list(maxfun = 100000))
)
drop1(morph_sur_mod, test = "Chi")



overdisp_fun(morph_sur_mod) #1.38


#diagnostic plots
simulateResiduals(fittedModel = morph_sur_mod, plot = TRUE)


testDispersion(simulateResiduals(fittedModel = morph_sur_mod))



summary(morph_sur_mod)


morph_sur$morph_prop_sur = morph_sur$`Second adult count` / morph_sur$`First adult count`


# Plot sex specific survival with predicted values and confidence intervals
fig_5 <- ggplot(morph_sur,
                aes(
                  x = morph_type,
                  y = morph_prop_sur,
                  shape = Pheromone,
                  colour = Pheromone,
                  fill = Pheromone
                )) +
  geom_point(size = 3, alpha = 0.3, position = position_dodge(width = 0.75)) +
```

```
  scale_color_manual(values = c("#1b9e77", "#d95f02"),
                     name = "Pheromone",
                     labels = c("Pheromone control", "Pheromone treatment")) +
  scale_fill_manual(values = c("#1b9e77", "#d95f02"),
                    name = "Pheromone",
                    labels = c("Pheromone control", "Pheromone treatment")) +
  scale_shape_manual(values = c(17, 16),
                     name = "Pheromone",
                     labels = c("Pheromone control", "Pheromone treatment")) +

  geom_boxplot(data = morph_sur,
               aes(x = morph_type, y = morph_prop_sur, fill = Pheromone),
               alpha = 0.4) +

  scale_x_discrete(labels = c("fig" = "Fighters", "scr" = "Scramblers")) +  # Rename x-axis
categories

  facet_grid(~Temperature) +
  labs(x = "Morph", y = "Late survival") +
  theme_light() +
  theme(panel.grid.minor = element_blank(),
        legend.key.size = unit(1, 'cm'),
        legend.key.height = unit(1, 'cm'),
        legend.key.width = unit(1, 'cm'),
        legend.title = element_text(size=14, face = "bold"),
        legend.text = element_text(size=14),
        legend.position = "bottom",
        axis.text = element_text(size = 14),
        axis.title = element_text(size = 14, face = "bold"),
        strip.text.x = element_text(size = 14, color = "black"),
```

```
        strip.text.y = element_text(size = 14, color = "black"),
        strip.background = element_rect(
          color = "black",
          fill = "#F2F4B5",
          size = 1.5,
          linetype = "solid"
        )
  )

# Display the graph
fig_5

ggsave(
  filename = "fig_5.pdf",
  plot = fig_5,
  dpi = 900
)



# Female proportion with trito only second count -------------------------
rm(list = ls(all = TRUE))
ls()

san_morph = read_excel("SS_heat_demography.xlsx", sheet = "morph")
san_morph$line = as.factor(san_morph$line)
san_morph$Pheromone = as.factor(san_morph$Pheromone)
san_morph$Temperature <- as.factor(san_morph$Temperature)
san_morph$fig = as.numeric(san_morph$fig)
san_morph$scr = as.numeric(san_morph$scr)
san_morph$female = as.numeric(san_morph$female)
```

```
san_morph$gen = as.numeric(san_morph$gen)

san_fem_prop = subset(san_morph, gen != 8 & count== "Second adult count")
san_fem_prop$male<- san_fem_prop$fig + san_fem_prop$scr

san_fem_prop$Temperature <- factor(san_fem_prop$Temperature, levels = c("Stable",
"Heatwave"))

fem_prop<-glmer(cbind(female, male) ~ (Temperature + Pheromone + as.numeric(gen))^2 +
(1|line), data = san_fem_prop, family = binomial,control = glmerControl(optimizer ='bobyqa',
optCtrl=list(maxfun=100000)))
drop1(fem_prop, test = "Chi")

fem_prop <- update(fem_prop,~.-Pheromone:as.numeric(gen))
drop1(fem_prop, test = "Chi")

fem_prop <- update(fem_prop,~.-Temperature:Pheromone)
drop1(fem_prop, test = "Chi")

fem_prop<-glmer(cbind(female, male) ~ Temperature + Pheromone + as.numeric(gen) + (1|line),
data = san_fem_prop, family = binomial,control = glmerControl(optimizer ='bobyqa',
optCtrl=list(maxfun=100000)))
drop1(fem_prop, test = "Chi")

overdisp_fun(fem_prop) #1.47

summary(fem_prop)

#diagnostic plots
simulateResiduals(fittedModel = fem_prop, plot = TRUE)
```

```
femprop_dummy <- data.frame(
  gen = rep(0:7, times = 4),  # 8 generations
  Temperature = rep(c("Stable", "Heatwave"), each = 16),
  Pheromone = rep(c("Control", "Treatment"), each = 8, times = 2)
)

predicted_femprop <- predict(
  fem_prop,
  newdata = femprop_dummy,
  re.form = NA,  # Exclude random effects
  type = "response",
  se.fit = TRUE
)


# Create predicted values dataframe
predicted_values_femprop <- data.frame(
  femprop_dummy,
  predict = predicted_femprop$fit,
  se = predicted_femprop$se.fit
)

# Add confidence intervals
predicted_values_femprop$lowerCI <- predicted_values_femprop$predict - 1.96 *
predicted_values_femprop$se
predicted_values_femprop$upperCI <- predicted_values_femprop$predict + 1.96 *
predicted_values_femprop$se

predicted_values_femprop$Temperature <- factor(predicted_values_femprop$Temperature,
levels = c("Stable", "Heatwave"))
```

```
fig_6a <- ggplot(san_fem_prop,  # Replace with your actual data frame
                 aes(
                   x = as.numeric(gen),
                   y = female / (male + female),
                   shape = Pheromone,
                   linetype = Pheromone,
                   colour = Pheromone,
                   fill = Pheromone
                 )) +
  geom_point(size = 2.2, alpha = 0.3, position = position_dodge(width =0.40)) +
  scale_x_continuous(breaks = 0:7) +

  # Add ribbon for confidence intervals
  geom_ribbon(data = predicted_values_femprop,
              aes(x = gen, ymin = lowerCI, ymax = upperCI, fill = Pheromone),
              alpha = 0.4, inherit.aes = FALSE) +

  # Add line for predicted values
  geom_line(data = predicted_values_femprop,
            aes(x = gen, y = predict, colour = Pheromone), linewidth = 1) +

  scale_color_manual(values = c("#1b9e77", "#d95f02"),
                     name = "Pheromone",
                     labels = c("Pheromone control", "Pheromone treatment")) +
  scale_fill_manual(values = c("#1b9e77", "#d95f02"),
                    name = "Pheromone",
                    labels = c("Pheromone control", "Pheromone treatment")) +
  scale_shape_manual(values = c(17, 16),
                     name = "Pheromone",
```

```
                    labels = c("Pheromone control", "Pheromone treatment")) +
scale_linetype_manual(values = c("solid", "dashed"),
                      name = "Pheromone",
                      labels = c("Pheromone control", "Pheromone treatment")) +

facet_grid(~Temperature) +

labs(
  x = "Generation",
  y = "Proportion of females"
) +

theme_light() +
theme(
  panel.grid.minor = element_blank(),
  legend.key.size = unit(1, 'cm'),
  legend.key.height = unit(1, 'cm'),
  legend.key.width = unit(1, 'cm'),
  legend.title = element_text(size=14, face = "bold"),
  legend.text = element_text(size=14),
  legend.position = "bottom",
  axis.text = element_text(size = 14),
  axis.title = element_text(size = 14, face = "bold"),
  strip.text.x = element_text(size = 14, color = "black"),
  strip.text.y = element_text(size = 14, color = "black"),
  strip.background = element_rect(
    color = "black",
    fill = "#F2F4B5",
    size = 1.5,
    linetype = "solid"
```

```
    )
  )

 # Display the graph
 fig_6a



 # female second count ----------------------------------------------------
 san_fem_first = subset(san_morph, count== "Second adult count" & total != 0 & gen != 8)
 hist(san_fem_first$female)


  san_fem_first$Temperature <- factor(san_fem_first$Temperature, levels = c("Stable",
"Heatwave"))


 fem_count1<-lmer(female ~ (Temperature + Pheromone + as.numeric(gen))^2  + (1|line), data =
san_fem_first)
 drop1(fem_count1, test = "Chi")


 fem_count1<-lmer(female ~ (Temperature + Pheromone + as.numeric(gen))^2  -
Temperature:as.numeric(gen)+  (1|line), data = san_fem_first)
 drop1(fem_count1, test = "Chi")


 fem_count1<-lmer(female ~ (Temperature + Pheromone + as.numeric(gen))^2  -
Temperature:as.numeric(gen) - Temperature:Pheromone +  (1|line), data = san_fem_first)
 drop1(fem_count1, test = "Chi")


 #diagnostic plots
 simulateResiduals(fittedModel = fem_count1, plot = TRUE)


summary(fem_count1)
anova(fem_count1)
```

```
fem_dummy <- data.frame(
  gen = rep(0:7, times = 4),  # 8 generations
  Temperature = rep(c("Stable", "Heatwave"), each = 16),
  #count = rep(c("First adult count", "Second adult count"), each = 16, times = ),
  Pheromone = rep(c("Control", "Treatment"), each = 8, times = 2)
)

predicted_fem <- predict(
  fem_count1,
  newdata = fem_dummy,
  re.form = NA,  # Exclude random effects
  type = "response",
  se.fit = TRUE
)


# Create predicted values dataframe
predicted_values_fem <- data.frame(
  fem_dummy,
  predict = predicted_fem$fit,
  se = predicted_fem$se.fit
)

# Add confidence intervals
predicted_values_fem$lowerCI <- predicted_values_fem$predict - 1.96 *
predicted_values_fem$se
predicted_values_fem$upperCI <- predicted_values_fem$predict + 1.96 *
predicted_values_fem$se
```

```
predicted_values_fem$Temperature <- factor(predicted_values_fem$Temperature, levels =
c("Stable", "Heatwave"))

fig_6b <- ggplot(san_fem_first,
                 aes(
                   x = as.numeric(gen),
                   y = female,
                   shape = Pheromone,  # Different shapes for points
                   linetype = Pheromone,  # Different line types for trends
                   colour = Pheromone,
                   fill = Pheromone
                 )) +
  geom_point(size = 2.2, alpha = 0.3, position = position_dodge(width =0.40)) +
  scale_x_continuous(breaks = 0:7) +

  scale_color_manual(values = c("#1b9e77", "#d95f02"),
                     name = "Pheromone",
                     labels = c("Pheromone control", "Pheromone treatment")) +
  scale_fill_manual(values = c("#1b9e77", "#d95f02"),
                    name = "Pheromone",
                    labels = c("Pheromone control", "Pheromone treatment")) +
  scale_shape_manual(values = c(17, 16),
                     name = "Pheromone",
                     labels = c("Pheromone control", "Pheromone treatment")) +
  scale_linetype_manual(values = c("solid", "dashed"),
                        name = "Pheromone",
                        labels = c("Pheromone control", "Pheromone treatment")) +

  # Add ribbon for confidence intervals
```

```
geom_ribbon(data = predicted_values_fem,
            aes(x = gen, ymin = lowerCI, ymax = upperCI, fill = Pheromone),
            alpha = 0.4, inherit.aes = FALSE) +

# Add line for predicted values
geom_line(data = predicted_values_fem,
          aes(x = gen, y = predict, colour = Pheromone, linetype = Pheromone), linewidth = 1) +

facet_grid(~Temperature) +
labs(
  x = "Generation",
  y = "Number of females"
)+
theme_light() +
theme(
  panel.grid.minor = element_blank(),
  legend.key.size = unit(1, 'cm'),
  legend.key.height = unit(1, 'cm'),
  legend.key.width = unit(1, 'cm'),
  legend.title = element_text(size=12, face = "bold"),
  legend.text = element_text(size=12),
  legend.position = "bottom",
  axis.text = element_text(size = 14),
  axis.title = element_text(size = 14, face = "bold"),
  strip.text.x = element_text(size = 13, color = "black"),
  strip.text.y = element_text(size = 13, color = "black"),
  strip.background = element_rect(
    color = "black",
    fill = "#F2F4B5",
    size = 1.5,
```

```
      linetype = "solid"
    )
  )

# Display the graph
fig_6b

fig_6<-ggarrange(
  fig_6a,
 fig_6b,
  nrow = 2,
  labels = c('a', 'b'),
  common.legend = TRUE,
  legend = "bottom",
  font.label = list(size = 20, face = "bold")
)
ggsave(
  filename = "fig_6.pdf",
  plot = fig_6,
  dpi = 900
)
```
